# Delta phase resets mediate non-rhythmic temporal prediction

**DOI:** 10.1101/643957

**Authors:** Jonathan Daume, Peng Wang, Alexander Maye, Dan Zhang, Andreas K. Engel

## Abstract

The phase of neural oscillatory activity aligns to the predicted onset of upcoming stimulation. Whether such phase alignments represent phase resets of underlying neural oscillations or just rhythmically evoked activity, and whether they can be observed in a rhythm-free visual context, however, remains unclear. Here, we recorded the magnetoencephalogram while participants were engaged in a temporal prediction task judging the visual or tactile reappearance of a uniformly moving stimulus. The prediction conditions were contrasted with a control condition to dissociate phase adjustments of neural oscillations from stimulus-driven activity. We observed stronger delta band inter-trial phase consistency (ITPC) in a network of sensory, parietal and frontal brain areas, but no power increase reflecting stimulus-driven or prediction-related processes. Delta ITPC further correlated with prediction performance in the cerebellum and visual cortex. Our results provide evidence that phase alignments of low-frequency neural oscillations underlie temporal predictions in a non-rhythmic visual and crossmodal context.

## Introduction

Neural oscillations reflect alternating states of higher or lower neural excitability, modulating the efficiency by which coupled neurons engage in mutual interactions (Buzsáki, 2006). As a result, neural communication and information processing has been shown to occur in a phase-dependent manner (Engel et al., 2001; Fries, 2005) reflected, for example, by fluctuations in perception thresholds correlating with the phase of ongoing oscillations (VanRullen, 2016). Based on these assumptions, oscillations were also linked to temporal predictions of upcoming relevant information (Arnal and Giraud, 2012; Engel et al., 2001; Rimmele et al., 2018). Studies have shown that animals can utilize predictive aspects of environmental stimuli in a way that reaction times are reduced (Gould et al., 2011; Lakatos et al., 2008; Rohenkohl and Nobre, 2011; Stefanics et al., 2010) or stimulus processing is enhanced (Cravo et al., 2013; Wilsch et al., 2015). By means of top-down induced phase resets of neural oscillations, phases of high excitability might be adjusted towards the expected onset of relevant upcoming stimulation in order to optimize relevant behavior (Schroeder and Lakatos, 2009).

Due to the rhythmic and therefore temporally highly predictable nature of many auditory stimuli such as speech or music, particularly in the auditory domain, many studies gathered evidence that oscillations reset and thereby adjust their phase towards rhythmic stimuli of various frequencies (Doelling and Poeppel, 2015; Giraud and Poeppel, 2012). Also in the visual domain, studies showed that neural oscillations align to temporal structure rhythmic visual input (Breska and Deouell, 2017b; Cravo et al., 2013; Gomez-Ramirez et al., 2011; Lakatos et al., 2008; Saleh et al., 2010). Other studies, however, reported a specific effect for visual temporal predictions only in the alpha band (8 – 12 Hz), although sensory input was provided at lower frequencies (Rohenkohl and Nobre, 2011; Samaha et al., 2015).

Moreover, whether temporal predictions indeed involve phase resets of endogenous neural oscillations remains a matter of debate (Breska and Deouell, 2017a; Doelling et al., 2019; Novembre and Iannetti, 2018). Despite their ecological relevance, using rhythms for the investigation of an involvement of oscillations in temporal predictions entails methodological and conceptual challenges. Rhythmic input leads to a continuous stream of regularly bottom-up evoked potentials, which are – at least – difficult to distinguish from top-down phase adjusted neural oscillations within the same frequency. Rather than by phase resets of endogenous neural oscillations, phase alignments across trials could therefore also be caused by stimulus-evoked potentials that just appear to be rhythmic during rhythmic stimulation (Doelling et al., 2019; Novembre and Iannetti, 2018; Zoefel et al., 2018).

Temporal prediction processes have further been shown to be reflected by slow buildups of neural activity, which ramps up until the predicted time point is reached; also called contingent negative variation (CNV; Breska and Deouell, 2017a; Macar et al., 1999). In a rhythmic temporal prediction context, such slowly ramping activity between subsequent stimulus pairs would also lead to significant phase-locking of low-frequency activity across trials, which again would be very difficult to be distinguished from phase-locking of phase-aligned endogenous neural oscillations. Conclusive evidence that temporal predictions involve phase resets of endogenous oscillations rather than stimulus-driven or prediction-evoked potentials is still lacking.

In addition, using only auditory rhythmic stimulation precludes the opportunity to link phase adjustments to a more general neural mechanism that predicts the temporal structure of any external input. If phase adjustments form the basis of tracking the temporal regularities of any relevant information, neural oscillations should align also to predictable temporal regularities that are inferred from input that does not itself comprise auditory rhythmic or discrete components, such as, for instance, uniform visual motion.

For this reason, we set out to investigate whether phase adjustments of neural activity can be observed for predictable visual motion stimuli. We measured magnetoencephalography (MEG) while healthy participants watched a visual stimulus continuously moving across the screen until it disappeared behind an occluder (Fig. 1A). We manipulated the time for the stimulus to reappear on the other side of the occluder. The task was to judge whether the stimulus reappeared too early or too late based on the speed of the stimulus earlier to disappearance. Hence, participants were required to temporally predict the correct time point of reappearance to be able to accomplish the task. Participants further performed a control task, in which the task was to judge the luminance of the reappearing stimulus instead of its timing. Importantly, physical appearance of both conditions was exactly the same in all aspects of the stimulation. Any purely stimulus-driven, bottom-up activity should therefore level out between the two conditions.

**Figure 1.**
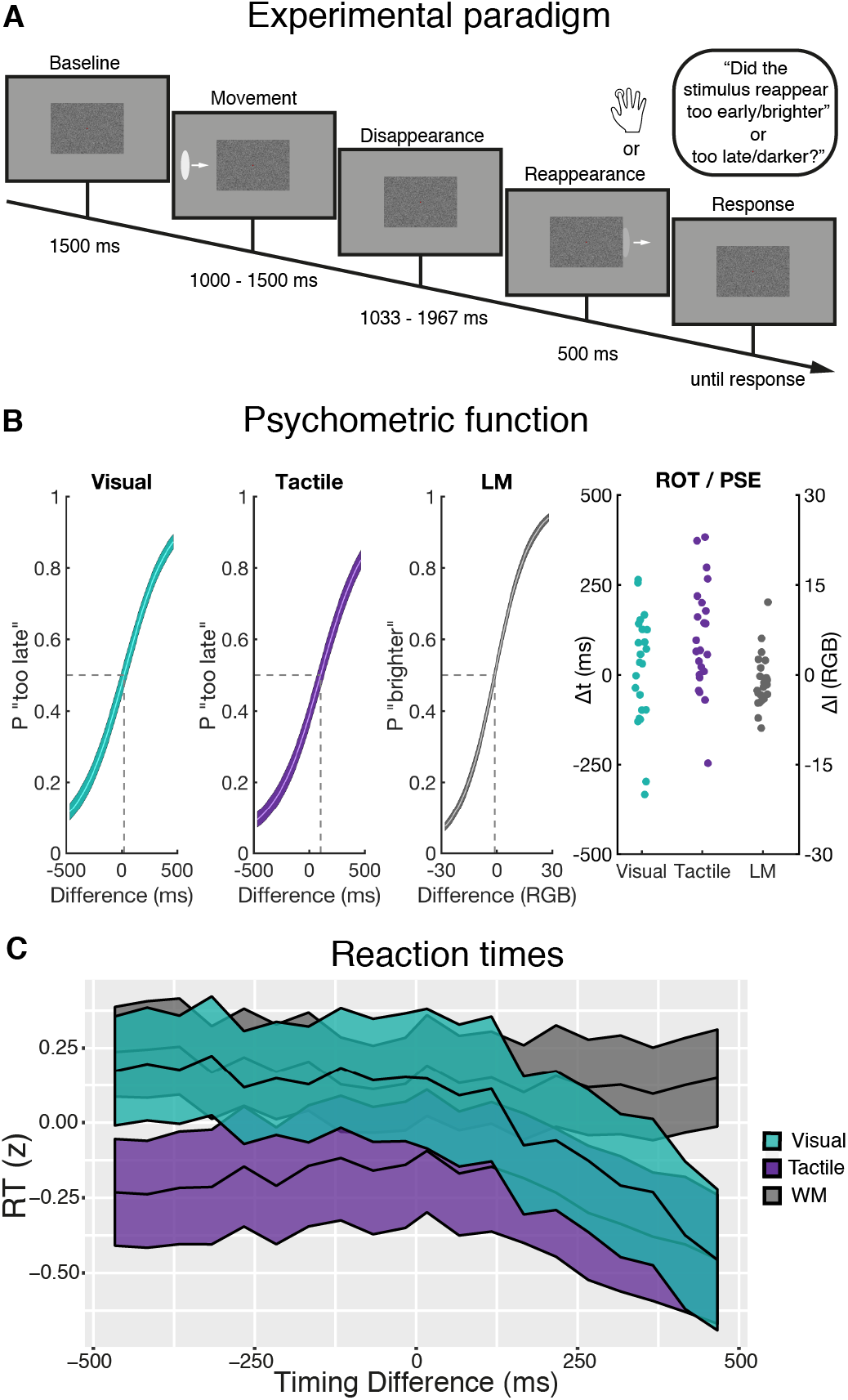
Experimental design and behavioral results. (**A**) A stimulus moved towards the center of the screen until it disappeared behind an occluder. The task was to judge whether the stimulus reappeared *too early* or *too late.* In the luminance matching condition, task was to judge whether the luminance became *brighter* or *darker.* Importantly, physical stimulation was exactly the same as in the visual prediction task. In the tactile temporal prediction task, at reappearance a tactile stimulus was presented contralateral to the disappearance of the visual stimulus. (**B**) Psychometric functions and individual ROT/PSE estimates. A timing difference of 0 refers to the objectively correct reappearance of the stimulus after 1,500 ms. Analogously, a luminance difference of 0 refers to equal luminance after reappearance provided in RGB values (see Methods). Colored areas depict standard errors of the mean (SEM). (**C**) Log-transformed and standardized reaction times for all timing differences (mean ± SEM). P = proportion; LM = luminance matching; t = time; l = luminance; RGB = red-green-blue.

Moreover, since it has been shown that sensory stimulation can lead to crossmodal phase adjustments also in relevant but unstimulated other modalities (Lakatos et al., 2007; Mercier et al., 2013), we further introduced a third condition in which a tactile instead of a visual stimulus was presented at reappearance. By contrasting it to the luminance matching control condition, we sought to determine whether phase alignments can be observed in regions associated with tactile stimulus processing, when sensory information was in fact only provided to the visual system.

We hypothesized that in the two temporal prediction tasks, as compared to the luminance matching control task, we would observe stronger inter-trial phase consistency (ITPC) within time windows between disappearance and expected reappearance. These phase alignments should particularly be observed at low frequencies, e.g., in the delta band, matching the temporal scale of the disappearance window (on average 1.5 s). Importantly, if such enhanced ITPC reflected phase resets of ongoing neural oscillations, we should not observe any changes in delta power during temporal predictions, as the amplitude of phase-resetting endogenous oscillations should not be altered. On the other hand, when stimulus-driven or prediction-evoked neural activity lead to the observed phase alignments, observed ITPC differences should be accompanied with differences in total delta power during temporal predictions, representing the evoked neural activity in each trial. Further, if the phase of neural oscillations indeed codes for the time point of the expected reappearance in each participant, participants showing a more consistent judgment of reappearance timing – as represented by a steep slope of the psychometric function – should have stronger ITPC during temporal predictions than participants who performed less accurately. If evoked neural activity underlies temporal predictions, these correlations should as well be accompanied by correlations between delta power and the steepness of the psychometric curve within the same brain region.

## Results

### Behavioral results

Participants did not receive feedback about the correctness of their response. This ensured that participants relied on their individual and subjective “right on time” (ROT) impression in the temporal prediction conditions and “point of subjective equivalence” (PSE) in the luminance matching condition. Across participants, there was no statistically significant bias towards “too early/darker” or “too late/brighter” responses in the visual temporal prediction (Δt (ROT_V_) = 13.15 ± 155.20 ms; *t*(22) = .41; *p* = .69; Cohen’s d = .09) or in the luminance matching task (ΔRGB (PSE) = −1.29 ± 4.54 RGB; *t*(22) = −1.36; *p* = .19; Cohen’s d = −.28), respectively (Fig. 1B). In the tactile temporal prediction task, participants showed a significant bias towards “too early” responses (Δt (ROT_T_) = 99.80 ± 150.00 ms; *t*(22) = 3.19; *p* = .004; Cohen’s d = .67).

To assess whether reaction times were dependent on the timing of the reappearing stimulus (Fig. 1C), we fitted a mixed-effect model to reaction times from all trials using the categorial variable *condition* (with the luminance matching task as reference level) and *timing difference* as well as their interaction as predictors. Since in the temporal prediction conditions we expected reaction times to be slowest for timing differences around zero and faster for high timing differences, we used a second-order polynomial term for *timing differences* (see Methods). Across all timing differences, reaction times were significantly faster in the tactile temporal prediction task as compared the luminance matching task (Δ = – 0.26; t = 14.21; *p* < 0.001), but not significantly different between the visual temporal prediction and the luminance matching task (Δ = −0.03; t = −1.34; *p* = 0.18). Across all conditions, reaction times linearly decreased with increase timing difference (Δ = −0.04; t = −s 2.48; *p* = 0.02) as well as showed a quadratic relationship with timing difference (Δ = 0.02; t = 2.42; *p* = 0.02). Importantly, as indicated by the interaction results, timing difference had a stronger negative linear (Δ = −0.13; t = −10.53; *p* < 0.001) and stronger negative quadratic influence on reaction times from the visual (Δ = −0.11; t = −8.18; *p* < 0.001) as well as a stronger negative quadratic influence on reactions times from the tactile temporal prediction task (Δ = −0.10; t = −7.52; *p* < 0.001) as compared to those from the luminance matching task (see Figure 1C and supplementary table S1 for the complete model output).

### Temporal prediction was associated with reduced beta power in sensory regions

Analyzing the neural data, we were first interested in investigating which frequency bands showed modulated spectral power during windows of temporal predictions in order to narrow down frequency bands of interest for further analyses. For that, we tested an average of spectral power across all sensors and conditions against a pre-stimulus baseline window. As a first step, we obtained a general overview of power modulations at each event in the experimental paradigm. Due to the jittered stimulation built into the design (see Materials and Methods), we computed cluster-based permutations statistics in three separate time windows (Fig. 2A) centered on: (a) the onset of the moving stimulus (“Movement”), (b) disappearance of the stimulus behind the occluder (“Disappearance”), and (c) reappearance of the stimulus (“Reappearance”).

**Figure 2.**
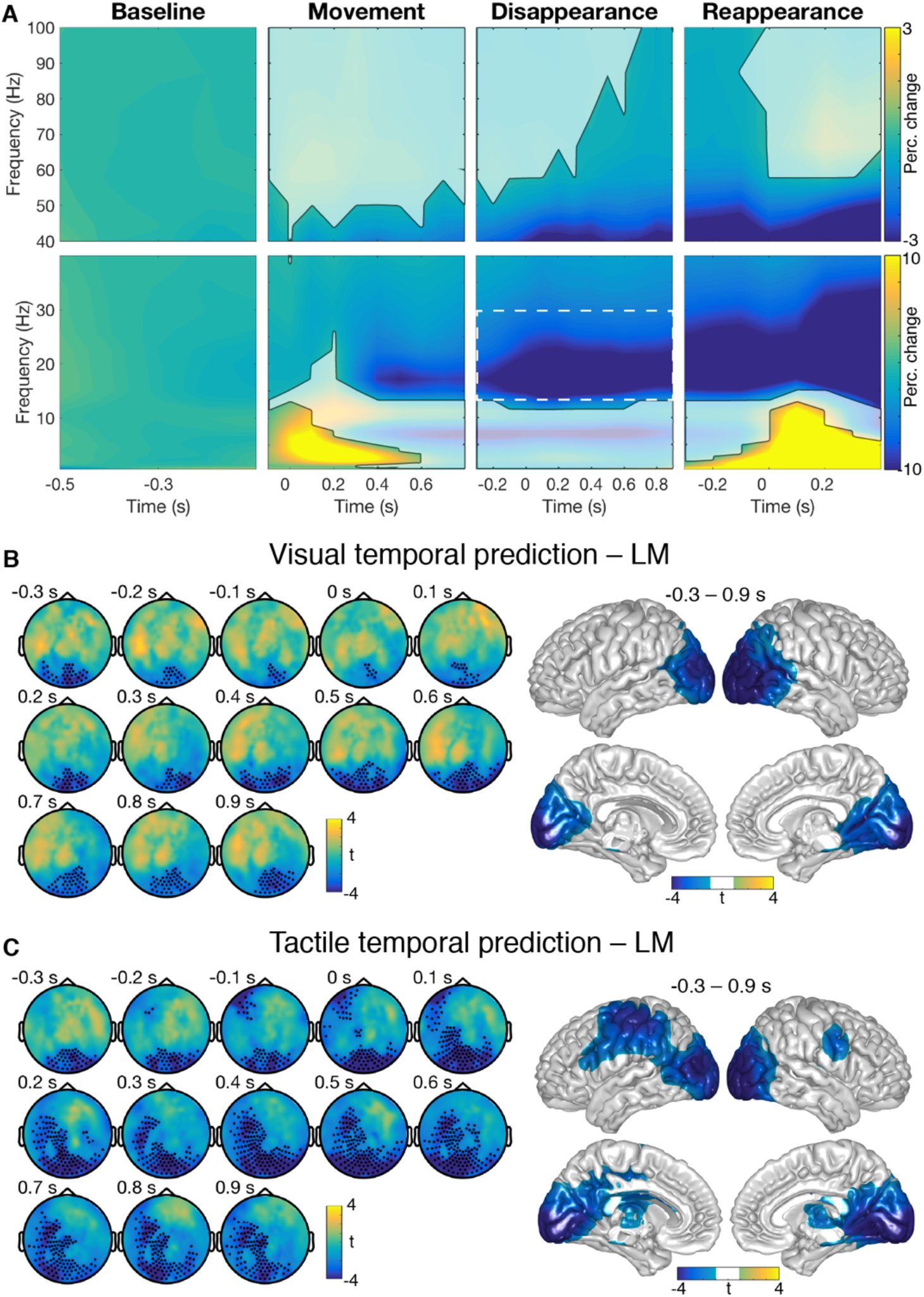
Power modulations during temporal prediction. (**A**) Spectral power averaged across sensors, conditions, and participants. Each window was centered on the different events within the paradigm and normalized with pre-stimulus baseline. Time 0 refers to the onset of each event. Cluster-based permutation statistics revealed significant power modulations as compared to baseline (unmasked colors). See also Fig. S1. (**B,C**) Difference between the two temporal prediction and the luminance matching task, respectively, within the beta band (13 – 30 Hz) in time bins around stimulus disappearance. Black dots indicate sensors of the clusters showing significant differences between the conditions. At source level, cluster-based permutation statistics revealed cluster of voxels showing significant differences between the conditions (colored voxels). LM = luminance matching.

In time bins around movement onset as well as reappearance of the stimulus, but not around disappearance, clusters of frequencies in the delta and theta range showed a statistically significant increase of spectral power as compared to the baseline window. All time windows further depicted a significant decrease of spectral power in frequencies within the beta and gamma range (all cluster *p*-values < .008). Importantly, even with using a liberal cluster alpha level of .05 (one-sided), we did not find a statistically significant modulation of delta power during the disappearance window. This was also not the case when reducing the test to sensors from occipital regions only (see Fig. S1).

Since we were most interested in examining modulations associated with temporal predictions, i.e., during the disappearance window, we further compared spectral power estimates between the temporal prediction tasks and the luminance matching task in all sensors within the disappearance window while ignoring the other windows. We restricted our analysis to the classical beta band ranging from 13 to 30 Hz, showing the strongest modulation as compared to baseline during the disappearance window. Cluster-based permutation statistics revealed reduced beta power during visual temporal prediction in occipital sensors during all time-bins of the disappearance window (cluster-*p* = .01). Source level statistics revealed a statistically significant decrease of beta power in a cluster of bilateral occipital voxels (cluster-*p* = .01). Beta power was further reduced during tactile prediction in a cluster of occipital as well as left lateralized frontocentral sensors (cluster-*p* = .002). At source level, a significant power reduction in the beta band was most strongly apparent in parts of bilateral visual as well as left-lateralized somatosensory cortex (cluster-*p* = .01).

### Delta inter-trial phase consistency was enhanced during temporal prediction

For the analysis of ITPC, we followed a similar approach. First, we tested ITPC differences to baseline in the three time windows for an average across all sensors and conditions using cluster-based permutation statistics. ITPC was significantly increased across a range of different frequencies in time bins around movement onset, disappearance and reappearance of the stimulus (all cluster-*p* < .001; Fig. 3A). For time windows centered on movement onset as well as reappearance significant ITPC increases were strongest in the delta to alpha range. At disappearance of the stimulus, significant ITPC increases were observed up to the low beta range with strongest increases in the delta band.

**Figure 3.**
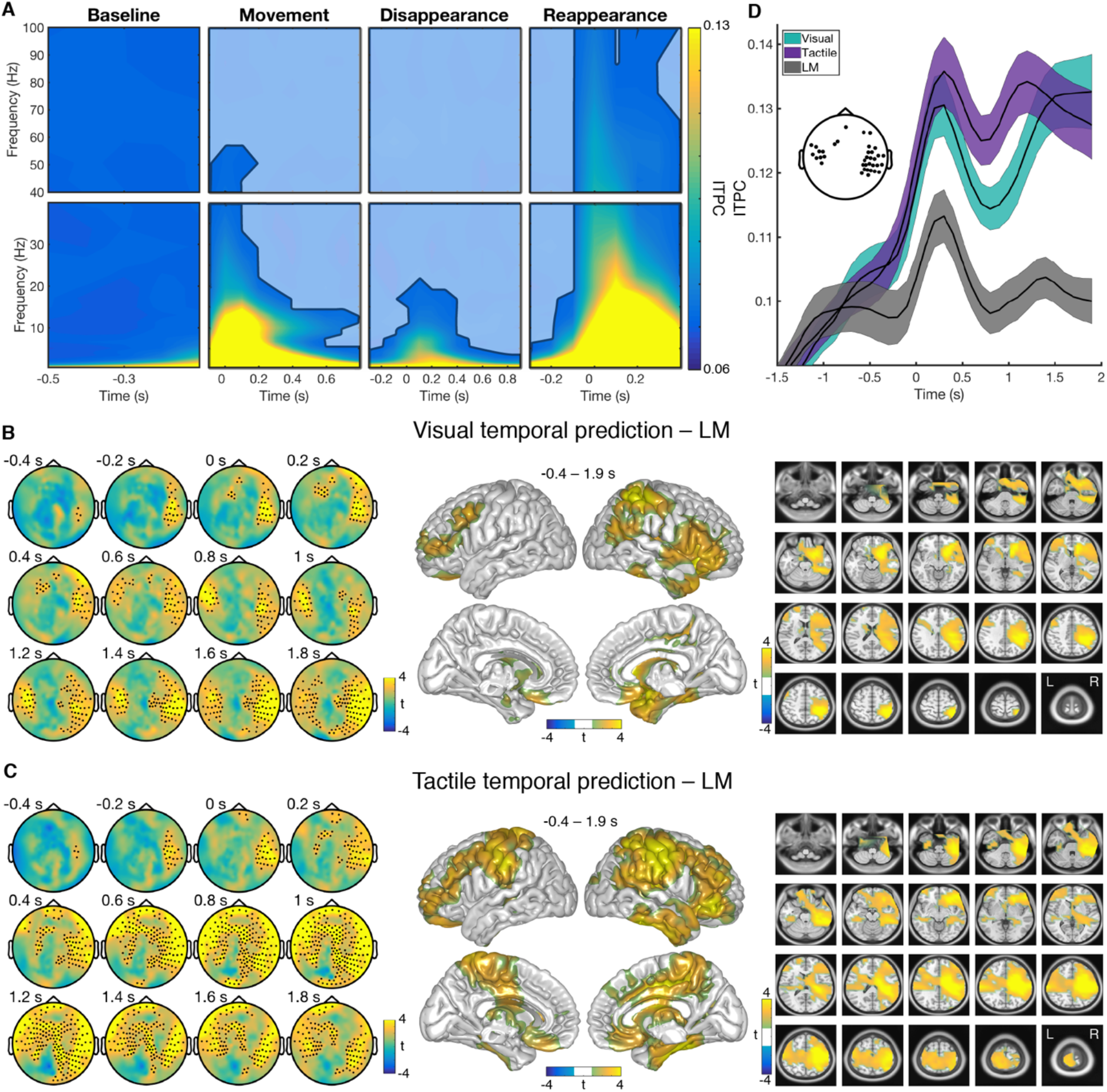
ITPC during temporal prediction as compared to luminance matching. (**A**) ITPC estimates averaged across sensors, conditions, and participants. Masked colors refer to non-significant ITPC modulations as compared to baseline (cluster-based permutation statistics). (**B,C**) Difference in ITPC between the visual or tactile prediction and the luminance matching task, respectively, within the delta band (0.5 – 3 Hz). For clarity, only every second time bin was plotted. Black dots indicate sensors of the clusters showing significant differences. On source level, clusters of voxels showing significant differences between the conditions are colored. See also Fig. S2 and S3 (**D**) Time course of absolute delta ITPC estimates within each condition for time bins centered around disappearance of the stimulus (time 0; mean ± SEM). ITPC estimates were averaged across channels that showed the top 20% of t-values for the comparison of the visual prediction with the luminance matching task (see topography). LM = luminance matching.

Hence, the delta band showed no increase in power but the strongest increase in ITPC as compared to baseline during the disappearance window for an average across all conditions (see Fig. 2A, 3A, and S1). For further statistical comparisons between conditions, we therefore restricted our analyses to an average of frequencies between 0.5 to 3 Hz (for condition-specific delta band ITPC differences to baseline during disappearance, see Fig. S2). For a better estimation of when differences in ITPC between the conditions became apparent, we enlarged the analysis of ITPC to time bins ranging from −1,900 ms to 1,900 ms centered on the disappearance of the stimulus. Note that in this enlarged analysis window the timing of the movement onset as well as the reappearance of the stimulus strongly jittered across trials. The effect of these events on ITPC estimates were thus strongly reduced (see Fig. S3).

We found two clusters that showed significantly stronger ITPC during visual temporal predictions as compared to luminance matching (Fig. 3B). One cluster included sensors from right temporal, frontal and occipital regions in time bins from −400 to 1,900 ms (cluster *p* < .001). The second cluster included left frontotemporal sensors in time bins ranging from 0 to 1,900 ms (cluster *p* = .01) Source level analysis revealed that for an average of the time window from −400 to 1,900 ms ITPC differences between the two conditions were strongest in right-lateralized central and inferior frontal voxels (cluster *p* < .001).

ITPC was also significantly enhanced in bilateral temporal sensors during tactile temporal predictions, evolving around −400 ms in right temporal sensors and shifting towards left hemisphere with ongoing disappearance time (cluster*p* < .001; Fig. 3C). In this contrast, however, differences in ITPC were more strongly apparent also in frontal and central sensors. Besides strongest differences in ITPC again in right superior parietal and inferior frontal voxels, source level analysis also revealed strong differences in bilateral somatosensory voxels for the contrast of tactile prediction to luminance matching (cluster*p* < .001).

Figure 3D depicts absolute delta ITPC estimates for all three conditions in the enlarged disappearance time window. Values were averaged across participants and all the sensors that exhibited the top 20% of t values in the ITPC contrast between visual temporal prediction and luminance matching between 0 and 1,500 ms (see Fig. 3B; similar results were obtained for sensors showing the top 10% or 5% of t values, see Fig. S3D). ITPC initially increased for all three conditions, but dropped down to stimulus movement level shortly afterwards in the luminance matching condition. ITPC in the visual as well as tactile temporal prediction tasks stayed elevated throughout the entire disappearance window.

### Control analyses on delta power differences between conditions

In contrast to ITPC, delta power did not significantly increase with disappearance of the stimulus in an average across conditions and channels as compared to baseline (see Fig. 2A and S1). Nevertheless, to examine whether channels showing the strongest differences in ITPC between conditions also show differences in delta power, we averaged delta power within the channels showing the strongest ITPC differences (same as in Fig. 3D) and compared power values from each of the two temporal prediction conditions with the luminance matching condition, respectively, within the same enlarged window of −1.900 to 1.900 ms around stimulus disappearance (Fig. 4).

**Figure 4.**
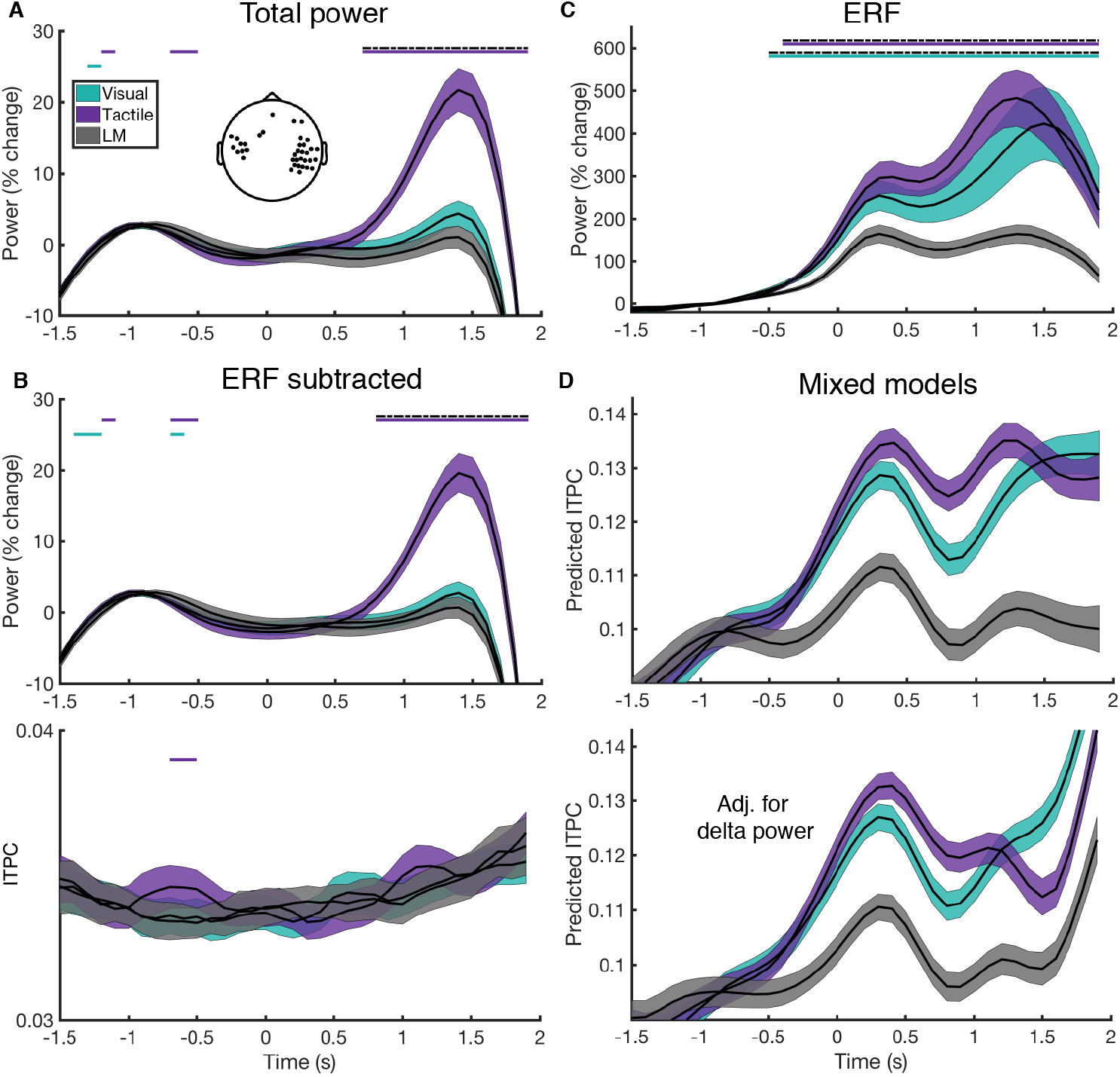
Delta power control analyses. (**A**) Time course of total delta power (0.5 – 3 Hz) in all conditions for the channels showing the strongest delta ITPC effect (same as in Fig. 3D). Time point 0 again refers to the complete disappearance of the stimulus. Colored lines depict uncorrected p-values below 0.05 from comparisons of the respective temporal prediction condition with the luminance matching task in each time bin. Dashed black lines depict p-values that survived the cluster-based permutation test. (**B**) The upper panel depicts the time course of induced delta power in each condition after a condition-wise subtraction of the ERF from each trial in the time domain. The lower panel depicts ITPC in each condition after ERF subtraction. (**C**) Delta power time course of the ERF, i.e., after averaging all trials in each condition in the time domain first. (**D**) Predicted delta ITPC values from mixed-effects regression models with an interaction term of *condition* and *time* as predictors for ITPC. Upper panel: without adjusting for delta power; lower panel: with adjusting for delta power by adding power as a covariate to the model. For a better comparability, standardized ITPC values were back-transformed to the original scale prior to plotting.

Figure 4A shows the time courses of total delta power in each condition. The overall pattern of the delta power time courses was largely different to the pattern of the ITPC time course in each of the conditions (compare to Fig 3D). In the visual temporal condition task as well as in the luminance matching task, delta power did not increase around disappearance of the stimulus and did not differ in any of the time bins between the two conditions during the disappearance window (even for uncorrected t-tests). Also in the tactile condition, delta power did not increase at around disappearance of the stimulus as observed for ITPC in this condition. However, it strongly increased in late time windows, showing significant differences in the tactile conditions as compared to the luminance matching task in time bins between 700 and 1900 ms after disappearance (cluster *p* < .001, dashed black line in Fig. 4A).

The combined pattern of an early delta ITPC increase and a late delta power increase in this condition could speak in favor of a CNV underlying the processes of temporal prediction. A CNV describes activity that is building up until the expected time point of an upcoming event is reached. After this time point, the build-up process sharply terminates (see, e.g. Breska and Deouell, 2017a; Macar et al., 1999). In such a scenario, ITPC would be increased as soon as the slow build-up process initiates (here at disappearance), but power increases might become observable only later in the prediction process. A phase-reset of ongoing oscillations, on the other hand, should not lead to an increase in delta power during the disappearance window.

To further investigate whether a CNV could explain the observed pattern of ITPC and power time courses, we computed additional control analyses. If a CNV would explain the increase in total power in the tactile condition, it should be locked to the disappearance of the stimulus and be present in each temporal prediction trial. Consequently, it should also be removed when subtracting the ERF from each trial in the time domain, before computing delta power in each single trial (i.e., when computing induced power). However, as the upper panel in Figure 4B shows, even after removing the ERF from each trial, delta power in the tactile condition was still strongly increased as compared to the luminance matching task in late disappearance time windows (cluster p < .001). Delta ITPC, on the other hand, was completely removed after subtracting the ERF (Fig. 4B lower panel).

Figure 4C depicts the delta power time course of the ERF itself in each condition. Delta power of the ERF increased for both, the tactile as well as the visual temporal prediction task, as compared to the control task. Similar to the ITPC effect, this increase already started in time bins shortly before disappearance (both cluster *p* < .001). Moreover, the strength of the increase in power in the visual temporal prediction task resembles the increase in the tactile task, and was not much stronger in the tactile task as observed for total power (Fig. 4A).

As a next step, we computed two mixed-effect regression models to examine the effect of delta power on ITCP. In one model, we used the variables *condition* and *time* as well as their interaction as predictors for ITPC only (Fig. 4D upper panel). In the other, we also added delta power as a covariate to the model in order to adjust for the variance explained by delta power (lower panel). After adding delta power as covariate, predicted ITPC values were reduced in the tactile prediction condition during late time windows of disappearance. However, they were still significantly different between both the visual and the tactile temporal prediction as compared to the luminance matching task, respectively, in all time bins during disappearance (see supplementary Table S2 for a complete model output).

### Delta ITPC, but not delta power, correlated to behavioral performance

We further hypothesized that if phase alignments of neural oscillations were indeed associated with temporal predictions, a participant who judged the reappearance of the stimulus within her individual subjectively correct ROT framework in a consistent manner should also exhibit stronger ITPC during temporal predictions, as a consistent timing judgement across trials should involve a similar phase across trials. The consistency of judgements can be inferred from the steepness of the psychometric function – the steeper the psychometric function, the more consistent the answers of the participant. We computed Pearson correlations of source level delta ITPC with the steepness of the psychometric function across participants and found statistically significant positive correlations in the visual (cluster *p* = .003) as well as in the tactile temporal prediction task (cluster *p* = .002; Fig. 5). Strongest correlations were found in the cerebellum and right lateralized early visual areas in both tasks. No clusters showing significant positive or negative correlations were observed in the luminance matching task (all cluster *p* > .1). If such correlations between phase alignments and behavior are related to evoked neural activity during temporal predictions, however, we should also observe similar correlation also between delta power and behavior. Hence, we averaged delta power within the voxels that showed the correlations between ITPC and behavior and computed Pearson correlations between this average and the steepness of the slope in each condition. We found no significant correlation between delta power and behavior in the visual (r = 0.31, *p* = 0.15) nor in the tactile temporal prediction task (r = 0.16, *p* = 0.47).

**Figure 5.**
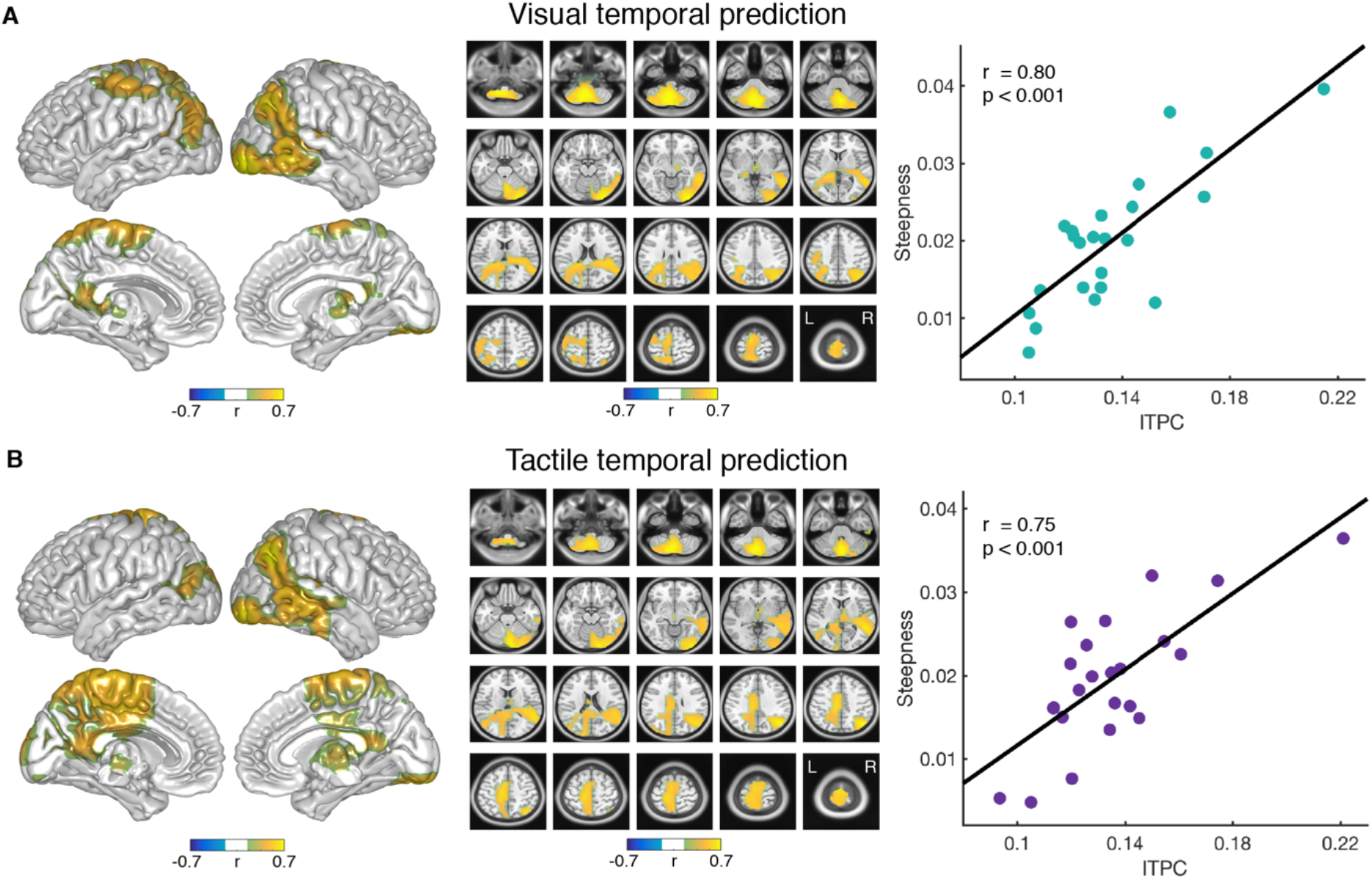
Correlation of ITPC to behavior. (A,B) Correlation of individual ITPC estimates with the individual steepness of the psychometric function within all voxels, shown in (A) for the visual prediction, and in (B) for the tactile prediction condition. ITPC estimates were averaged within the delta band and time windows of 0 to 1,000 ms centered on the disappearance of the stimulus. Only the clusters of voxels showing significant correlations are colored. In the scatter plots, each dot represents one participant and ITPC estimates were averaged across all voxels within the clusters of significant correlations. There was no significant correlation observed for the luminance matching condition or between delta power and behavior.

### ITPC did not correlate with eye movements

A potential confound for the observed effects in ITPC could be that participants tracked the moving stimulus with their eyes to be able to judge the correct time point of reappearance. Thus, consistent horizontal eye movements with the speed of the stimulus might lead to enhanced ITPC in the delta band. To make sure that differences in eye movements do not explain the observed differences in ITPC between the conditions, we analyzed horizontal eye movements recorded by an eye tracker (ET) during the MEG measurement. Figure 6A depicts condition-wise horizontal eye positions averaged across all participants and centered on the disappearance of the stimulus, showing no systematic differences between the conditions.

Moreover, if horizontal eye movements would explain the effects in ITPC, we should observe the same effects between the conditions when we compute ITPC for the ET data. Differences in ITPC between the two temporal prediction conditions and the luminance matching condition are depicted Figure 6B and C. Using cluster-based permutation statistics, we did not observe any time-frequency cluster that revealed significant differences between the conditions (all cluster *p* > .1).

**Figure 6.**
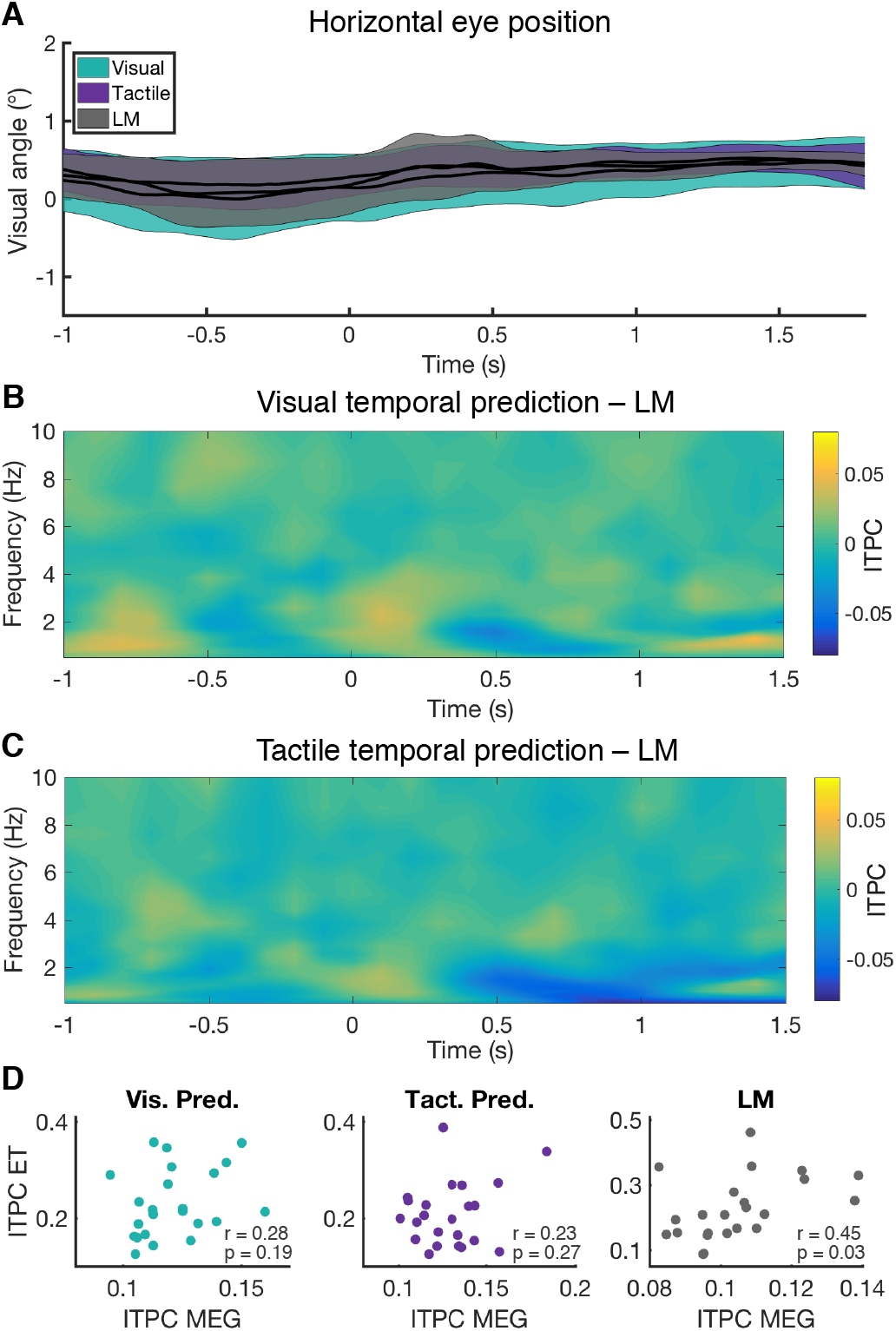
Analysis of horizontal eye movements. (**A**) Condition-wise eye positions centered on stimulus disappearance (time 0 s) and averaged across all participants. A visual angle of 0° refers to the fixation dot (1° visual angle roughly corresponds to 1 cm on the screen). Colored areas depict SEM. (**B,C**) Differences in ITPC between (**B**) the visual prediction and (**C**) the tactile prediction condition to the luminance matching condition in low frequencies and time bins around disappearance of the stimulus (time 0 s). Utilizing cluster-permutation statistics, no clusters of significant differences were observed between the conditions. (**D**) Condition-wise correlations between ITPC estimates obtained from the eye tracker data and the MEG sensors across all participants. ET = eye tracker.

Further, we tested whether there are any significant correlations between individual ITPC values obtained from the MEG data and from the ET data. We averaged ITPC values from a time window of 0 to 1.000 ms and again used the top 20% of channels showing the strongest effect for ITPC for the MEG data (for channels see Fig. 3D), and did not observe significant correlations between the ITPC values obtained from MEG and ET data in the temporal prediction tasks (Fig. 6D). The strongest correlation was found in the luminance matching condition, which suggests that the ITPC differences found in the MEG data cannot be explained by horizontal eye movements during temporal predictions.

## Discussion

Our task design enabled us to disentangle phase resets of ongoing neural oscillations from evoked event-related potentials. We found that phase alignments, but not power, were stronger in the context of temporal predictions than in a task where temporal structure was less relevant. This supports the hypothesis that phase adjustments of ongoing neural oscillations, and not stimulus-driven or prediction-evoked activity, form the neuronal basis of temporal prediction processes and suggest that this framework can be extended to predictions that have to be inferred from stimulation that does not itself comprise rhythmic and discrete components. The strength of the observed phase adjustments further correlated with the ability to consistently judge the temporal reappearance of the stimulus across participants, suggesting also a functional relevance of the observed phase adjustments for temporal predictions.

### Cross-modal temporal predictions are reflected by a beta power reduction in both sensory systems

It has been suggested that temporal predictions of upcoming events might be mediated by neuronal oscillations in the delta and beta frequency range (Arnal and Giraud, 2012). The enhanced phase consistency of delta oscillations as well as the power modulations in the beta band observed in the current study are in line with this hypothesis. However, earlier reports on beta power modulations during temporal predictions are inconsistent. On the one hand, studies found that beta power was even increased shortly before the onset of the expected stimulus in auditory (Arnal et al., 2015) and visual rhythmic stimulation (Saleh et al., 2010). On the other hand, van Ede et al. (van Ede et al., 2011) found that predicting the onset of a tactile stimulus was specifically associated with a reduction of beta power in contralateral tactile areas and accompanied by faster reaction times. The authors suggest that a reduction in beta power might signal preparatory processes in the sensory system that expects the upcoming event.

The observed decrease in beta power in task-relevant sensory regions in the current study largely match the results reported by van Ede et al. (van Ede et al., 2011). During visual temporal predictions, beta band power was reduced in visual sensory regions as compared to the visual control condition during the entire disappearance time. During crossmodal predictions, in which temporal information was provided to the visual system, but reappearance was expected in the tactile domain, beta band power was decreased in both, visual as well as tactile regions.

Since also in the luminance matching condition participants expected to perceive a visual stimulus, preparatory processes alone cannot explain this reduction in beta power. This is especially the case in the crossmodal condition, in which no visual stimulus was expected, but stronger decreases in beta were also observed in visual areas. Moreover, since we observed beta decreases also in tactile regions at the time of visual stimulus disappearance, the decrease could not solely be an effect of external stimulation.

Beta decreases observed during temporal predictions might therefore relate to more than only preparatory processes to an upcoming stimulus. Cross-modal decreases in beta band activity in both the temporal information providing visual as well as the stimulation expecting tactile system might reflect that both sensory modalities are continuously involved in temporal prediction processes, not only in processes preparing for the upcoming stimulation. We found no significant increases in beta power during temporal predictions. Whether decreases in beta power are associated with non-rhythmic temporal predictions while increases might reflect temporal predictions during rhythmic stimulation, remains subject to future research.

### Enhanced ITPC cannot be explained by event-related increases in neural activity

In earlier investigations of phase adjustments to external predictive stimulation, participants were mostly presented with streams of auditory rhythmic input. Rhythmic and discrete input, however, also causes strongly evoked brain activity within the same frequency range, which makes it difficult to disentangle streams of evoked activity from entrained endogenous neural oscillations (Novembre and Iannetti, 2018; Zoefel et al., 2018). Our results provide evidence that phase alignments of low-frequency fluctuations observed during temporal predictions cannot solely be explained by stimulus-driven, bottom-up evoked brain activity (see also, Doelling et al., 2019; Kösem et al., 2018). In the current study, we aimed at reducing stimulus-evoked brain responses to a minimum by presenting participants with a continuously moving stimulus instead of several discrete stimuli. We were particularly interested in the time point at which the stimulus transiently disappeared behind an occluder (as opposed to sharp onsets and offsets in discrete rhythmic stimulation). At disappearance, we did not observe an increase in low-frequency power as compared to pre-stimulus baseline in any of the conditions, which could explain an increase of phase alignments after disappearance of the stimulus. Moreover, by using an experimental design in which physical stimulation at disappearance was exactly the same during temporal predictions as well as the control condition, we controlled for brain responses that could have been driven by bottom-up, stimulus-processing activity and would therefore not be specific to temporal predictions. Importantly, delta ITPC, but not power, was stronger during temporal predictions at and after disappearance of the stimulus, suggesting that delta phase alignments during temporal predictions cannot be solely related to brain responses evoked by the offset of the visual movement.

It has been further suggested that a CNV, i.e., activity that is ramping up until the expected time point is reached, might underlie enhanced phase alignments during temporal predictions (Breska and Deouell, 2017a). CNVs have often been observed in timing tasks (e.g., Macar et al., 1999; Pfeuty et al., 2003; Praamstra et al., 2006), and such ramping activity initialized by temporal predictions would, besides an increase in power, also lead to increased phase alignments as reflected by enhanced ITPC during temporal predictions. These increases in activity are therefore not caused by the physical stimulation itself but specifically related to temporal predictions. As described above, the observed pattern of an early delta ITPC increase and a late delta power increase in our tactile prediction condition could speak in favor of a CNV underlying the processes of temporal prediction (see Fig. 3D and 4A). However, there are several aspects that argue against an involvement of a CNV in our data.

First of all, if in our data a CNV underlay temporal predictions, we should have observed a late power increase also in the visual temporal prediction task, in which participants also focused temporal predictions but saw the exact same physical stimulation as in the control task. Even using uncorrected t-tests, however, we did not observe total delta power differences between the two conditions in any of the time bins after stimulus disappearance. Since we see strong ITPC increases in both temporal prediction condition, but a delta power increase only in the tactile condition, it is unlikely that CNV-like activity would explain the phase alignments observed in *both* temporal predictions tasks.

Moreover, by subtracting the ERF from each single trial, all activity that is phase-locked to the disappearance of the stimulus should be removed from the data, that is, all activity related to a phase-reset of oscillations *as well as* all activity reflecting event-related potentials. However, also after subtracting the ERF, the strong delta increase in the tactile condition was still observable. This suggests that the increase in power in the tactile condition was not associated to the temporal prediction processes locked to the disappearance of the stimulus. Since the reappearance of the stimulus strongly jittered in relation to the time point of disappearance and the tactile condition was the only condition in which a sharp-onsetting tactile stimulus was presented, it is likely that this delta power increase in late windows was caused by the presentation of the tactile stimulus. In contrast to power, however, delta ITPC was completely removed after subtracting the ERF, which confirms that subtracting the ERF reliably removed all disappearance-locked activity.

Further, as stated above, averaging across all trials, i.e., computing the ERF, would capture all activity from each trial which is locked to disappearance of the stimulus, i.e., phase-reset oscillatory activity and/or event-related potentials. In contrast, unlocked activity should be removed by the averaging. If a CNV caused the late power increase in the tactile condition, this pattern of a late increase in power should also be observable for the power time course of the ERF. A phase-reset of oscillatory activity, on the other hand, would rather cause an ERF power time course that shows differences already at early time windows of the disappearance, as is the case in our data. The time course of delta power of the ERFs, therefore, speak against a CNV representing the power increase but, rather, for oscillatory activity that resets its phase at disappearance.

Therefore, instead of a CNV causing phase alignments of slow fluctuations across trials (as described above) the opposite might hold, i.e., phase resets of oscillatory activity might actually, after averaging, lead to results erroneously suggesting a CNV. If so, studies that observed a CNV after averaging, could have in fact also extracted all oscillatory activity that has reset its phase after the onset of a temporal cue. Only if a CNV was present in single trial data, and not only after averaging, such event-related slow fluctuations would indeed relate to single trial temporal predictions. In our data, however, we did not observe a temporal prediction related increase in *total* delta power, which is computed on single trial time courses *before* averaging. An increase in power was only visible *after* averaging all trials in the time domain first. Thus, the lack of a power increase in total power together with a CNV-like power increase after averaging across trials suggests that neural oscillations reset their phase according to the temporal structure of the stimulation, but did not alter in amplitude on a single trials basis.

Taken together, we observed strong ITPC differences between the conditions but no (total) power differences that could be explained by event-related potentials such as a CNV. Instead of evoked or CNV-like activity, our results therefore suggest that the phase alignments observed during temporal predictions are associated to neural oscillations that adjusted their phase to the temporal structure of the stimulation in order to predict the reappearance of the upcoming stimulation.

### Neural oscillations at low frequencies adapt to the temporal structure of non-rhythmic visual motion stimulation

Earlier studies have observed that neural oscillations entrain towards rhythmic sensory input to track the low-frequency temporal regularities of the stimulation, especially in the auditory domain (Giraud and Poeppel, 2012). Such phase entrainment does not only occur in the delta band but can flexibly adapt to the frequency of the external input also at higher frequencies such as the theta or the alpha band during auditory stimulation (Doelling and Poeppel, 2015). However, in the visual system, evidence for the tracking of temporally predictive input by neural oscillations is not as clear. On the one hand, studies showed that the phase of neural oscillations is involved in temporal predictions of low-frequency visual input (Breska and Deouell, 2017a; Cravo et al., 2013; Saleh et al., 2010; Wilsch et al., 2015). On the other hand, studies suggested that temporal predictions in the visual system were specific to the alpha band, although sensory input was provided at lower frequencies (Rohenkohl and Nobre, 2011; Samaha et al., 2015). Rohenkohl and Nobre (Rohenkohl and Nobre, 2011), for instance, used rhythmically presented visual stimuli at 2.5 and 1.25 Hz moving across the screen until it disappeared behind an occluder. Nevertheless, neural oscillations exclusively from the alpha band showed modulated activity associated with temporal predictions during the disappearance time. They found no phase locking of oscillations in lower frequencies.

In the current study, we provide further evidence that neural oscillations from the delta band show enhanced phase alignment during visual temporal predictions across trials. In order to adapt to the temporal regularity of the presented visual stimulus, delta frequencies in a wide network of parietal and frontal brain areas exerted more consistent phase resets at around the time point of disappearance of a monotonically moving stimulus as compared to a luminance matching control condition. The strength of this phase adjustment in each participant correlated with the consistency in judging a reappearance of the visual stimulus as too early or too late. This was the case only in the temporal prediction tasks, which underlines the behavioral relevance of the observed phase adjustments for temporal predictions.

Importantly, our study suggests that the mechanism of phase adjustments for temporal predictions can be extended to external stimulation that does not as such involve rhythmic or discontinuous stimulation. We found that low-frequency oscillations can adjust their phase also to the temporal structure of external stimulation that had to be inferred from uniform visual motion. This is also in line with recent studies reporting enhanced performance as well as an involvement of delta phase for non-rhythmic, yet predictable stimulation in the auditory (Herbst and Obleser, 2019) as well as the visual domain (Breska and Deouell, 2017a; but see Obleser et al., 2017 and Breska and Deouell, 2017b for a discussion about the rhytmicity of their non-rhythmic visual stimulation). While both studies involve onsets of discrete stimuli, they show that delta phase was involved in temporal prediction processes during stimulation that was not itself purely rhythmic. By showing that the phase of neural oscillations also align to a rhythm-free, non-discrete, unimodal visual as well as crossmodal visuotactile stimulation, our results further indicate that the framework of phase adjustments during temporal predictions might be generalized also to other, if not all, forms of temporally predictive external stimulation.

### Phase resets occurred in a network of frontoparietal and sensory brain areas

We observed enhanced ITPC values in a network of mostly frontal and parietal brain areas during visual as well as crossmodal temporal predictions. Similarly, Besle et al. (2011) observed significant phase entrainment to audiovisual stimulation in a wide network of distributed areas including parietal and inferior frontal areas. These observations support the notion that brain areas involved in temporal predictions may constitute a frontoparietal timing network (Coull and Nobre, 2008; Rimmele et al., 2018).

Further, we found enhanced ITPC values also in early somatosensory areas contralateral to the disappearance of the purely visual stimulus during crossmodal temporal predictions, despite the fact that prediction-relevant information was provided only by a moving visual stimulus. This supports evidence reported earlier showing that stimulation within one modality can crossmodally reset the phase of ongoing low-frequency in other modalities, which might be an important mechanism for multisensory integration processes (Lakatos et al., 2007; Mercier et al., 2013).

Similarly, we expected to find enhanced ITPC during temporal predictions in early visual areas. In fact, increased delta ITPC as compared to baseline were also observed in occipital sensors (see Fig. S2), but they were not significantly different between the conditions. However, we found that voxels in early visual areas showed strong correlations between individual ITPC estimates and the steepness of the psychometric function in both temporal prediction tasks, but not in the luminance matching task. This suggests that consistent phase resets of delta oscillations within visual areas might have supported consistent timing judgments with the participants’ subjective timing frameworks. This indicates an involvement also of the visual system in processes related to temporal prediction.

Moreover, strong correlations between ITPC and behavior were also observed in the cerebellum, supporting earlier reports on a involvement of the cerebellum in temporal prediction processes (Breska and Ivry, 2016). Roth and coworkers (Roth et al., 2013), for instance, found that cerebellar patients were significantly impaired in recalibrating sensory temporal predictions of a reappearing visual stimulus. This finding is of particular interest as we adapted the authors’ experimental paradigm for the use in the current study. Theirs and our results therefore indicate that the cerebellum might be crucially involved in accurate and consistent judgments of temporal regularities deployed in perceiving object motion.

## Conclusions

We provide evidence that the phase of neural oscillations can adjust to the temporal regularities of external stimulation and do not arise as a byproduct of stimulus-driven or prediction-related evoked potentials. Such phase alignments could provide a key mechanism that predicts the onset of upcoming events in order to optimize processing of relevant information and thereby adapt behavior. We show that temporal information provided to one modality leads to phase adjustments in another modality when crossmodal temporal predictions are necessary, providing further evidence that such crossmodal phase resets could be the neuronal basis of multisensory integration processes. Moreover, phase alignments were observed for unimodal visual as well as crossmodal visuotactile non-rhythmic and non-discrete stimulation, suggesting a generalizability of phase resets as a mechanism for temporal predictions to all forms of external stimulation. Taken together, our results provide important further insights into the neural mechanisms that might be utilized by the brain to predict the temporal onsets of upcoming events.

## Materials and Methods

### Participants

Twenty-three healthy volunteers (mean age ± standard deviation (SD): 27.13 ± 4.30 years; 20 females; all right-handed) took part in the study. They gave informed written consent and were monetarily compensated with 13 €/hour for participation. All volunteers had normal or corrected-to-normal vision, normal touch, as well as no background of psychiatric or neurological disorder. The ethics committee of the Medical Association Hamburg approved the study protocol (PV5073), and the experiment was carried out in accordance with the approved guidelines and regulations.

### Experimental procedure

The experimental paradigm used in the current study was adopted from an earlier report investigating visual temporal predictions in cerebellar patients (Roth et al., 2013). Our experiment consisted of three conditions: a *visual* temporal prediction task, a crossmodal *(tactile)* temporal prediction task, and a *luminance matching* (control) task. The trials of all conditions started with the presentation of a randomly generated, white noise occluder (size: 7.5° x 11.3° (h x w)) that was smoothed with a Gaussian filter (imgaussfilt.m in MATLAB) and presented in the middle of the screen against a grey background screen (luminance: 44 cd/m^2^; corresponds to 115 red-green-blue (RGB) values in our setting; see Figure 1A). At the center of the occluder, a red fixation dot was presented. We instructed participants to fixate this dot throughout the entire trial. After 1500 ms, an oval stimulus (size: 3.5° x 1.0°) set on in the periphery of the screen, moving towards the occluder with a speed of 6.9 °/s. The luminance of the stimulus differed in all trials between 120 to 161 cd/m^2^ (6 steps, counterbalanced, corresponds to 170 to 220 RGB). For half of the participants, the stimulus started on the left side of the occluder and moved from left side towards the right side. For the other half, the stimulus started on the right side and moved from right to left. The direction of movement was kept constant for each participant throughout the entire experiment. In each trial, the starting point of the stimulus differed such that the stimulus took 1,000 to 1,500 ms to disappear completely behind the occluder from starting point, randomly jittered with 100 ms (counterbalanced). The size of the occluder and the speed of the stimulus were chosen so that the stimulus would need exactly 1,500 ms to reappear on the other side of the occluder. However, we manipulated the timing and the luminance of the reappearing stimulus. In each trial, the reappearance of the stimulus differed between ±17 to ±467 ms (randomly jittered, but counterbalanced in steps of 50 ms; corresponds to ±1 to ±28 frames with a jitter of 3 frames at 60 Hz) from the correct reappearance time of 1,500 ms. Hence, the stimulus was covered by the occluder for 1,033 to 1,967 ms and was reappearing at 20 different time points. In the visual prediction task as well as in the luminance matching task, we also manipulated the luminance of the reappearing stimulus relative the luminance the stimulus had before disappearance in each trial (jittered, but counterbalanced between ±1 to ±40 cd/m^2^, also using 20 different values; corresponds to ±1 to ± 28 RGB in steps of 3 RGB to make it similar to the timing manipulation). After reappearance, the stimulus moved to the other side of the screen for 500 ms with the same speed until it set off the screen. The occluder was presented throughout the entire trial.

By manipulating the timing as well as the luminance in both conditions, we made sure that both, the visual temporal prediction as well as the luminance matching task had the exact equal physical appearance throughout all trials. They only differed in their cognitive set. In the visual temporal prediction task, we asked participants to judge whether the stimulus was reappearing *too early* or *too late* based on the speed the stimulus had earlier to the occluder (which was kept constant throughout the entire experiment). In the luminance matching task, participants were asked to judge whether the luminance of the reappearing visual stimulus became *brighter* or *darker* as compared to the stimulus earlier to disappearance. Participants answered by pressing one of two buttons with their index or middle finger of the hand contralateral to the reappearing stimulus.

The tactile temporal prediction task was equal to the visual temporal prediction task, with the only difference that a tactile stimulus instead of a visual was presented at the time of reappearance to the right or left index finger (depending on which side the stimulus was expected to reappear behind the occluder). The tactile stimulus was presented by means of a Braille piezostimulator (QuaeroSys, Stuttgart, Germany; 2 x 4 pins, each 1 mm in diameter with a spacing of 2.5 mm), pushing up all eight pins for 200 ms. At that time, nothing happened on the screen. Participants gave their answer with the same hand as in the other two conditions (i.e., with the hand that was not stimulated by the Braille stimulator). Response mapping of the two buttons was counterbalanced across all participants. As soon as participants gave their answer, the fixation dot turned dark grey for 100 ms to indicate that the response was registered. However, participants did not receive trial-wise feedback about the correctness of their response. After a short delay of 200 ms, the white-noise occluder was randomly re-shuffled to signal the start of a new trial.

All three conditions were presented block-wise. At the beginning of each block, participants were informed about the current task. The order of presentation of the conditions was kept constant for each participant, but was randomized across participants (counterbalanced). At the end of each block, they were informed about the overall accuracy of their answers within the last block and were allowed to rest as long as they wanted. Each participant performed two sessions at two different recording days. The experimental procedure was kept constant across both sessions, i.e., movement direction, response mapping, as well as condition order did not change in the second session for individual participants. Each session comprised twelve blocks, i.e., four blocks per condition. Each block consisted of 60 trials, resulting in a total number of 480 trials per condition or 1,440 trials in total. Due to technical difficulties, for one participant we only acquired data from one session with a total number of 720 trials.

At the beginning of each recording day, participants performed a short training of all conditions to get familiar with the overall experimental procedure and the stimulus material. This training took part in the same environment as the subsequent recording session. At the end of the second recording day, participants filled a questionnaire asking for any specific strategy they might have used for the temporal prediction task.

We used MATLAB R2014b (MathWorks, Natick, USA; RRID: SCR_001622) and Psychtoolbox (Brainard, 1997) (RRID: SCR_002881) on a Dell Precision T5500 with Ubuntu 64-bit operating system (Version: 16.04.5 LTS) for stimulus presentation. The visual stimuli were projected onto a matte backprojection screen at 60 Hz with a resolution of 1,920 × 1,080 pixels positioned 65 cm in front of participants. To mask the sound of the Braille stimulator during tactile stimulation, we presented participants with auditory pink noise at sampling rate of 48 kHz and volume of 85 dB using MEG-compatible in-ear headphones (SRM-252S, STAX Limited, Fujimi, Japan) during all experimental blocks.

### Data acquisition and pre-processing

MEG was recorded at a sampling rate of 1,200 Hz using a 275-channel whole-head system (CTF MEG International Services LP, Coquitlam, Canada) situated in a dimly lit, sound attenuated and magnetically shielded chamber. We additionally recorded electrical eye, muscle and cardiac activity with Ag/AgCl-electrodes in order to have a better estimate for endogenous artefacts. Online head localizations (Stolk et al., 2013) were used to navigate participants back to their original head position prior to the onset of a new experimental block if their movements exceeded five mm from their initial position. The initial head position from the first recording day was saved so that participants could be navigated back to their initial head position also during the second recording day. This assured comparable head positions of each participant across sessions. Five malfunctioning channels were either not recorded or excluded from further analysis for all participants. To further control for eye movement artifacts, eye movements were tracked with an MEG-compatible EyeLink 1000 Long Range Mount system (SR Research, Osgoode, Canada).

We analyzed reaction time data using R (R Core Team, 2014) (RRID: SCR_001905) and RStudio (RStudio Inc., Boston, USA; RRID: SCR_000432). Trials with reaction times longer than three standard deviations were excluded from analysis. Due to the right-skewed nature of reaction times, reaction time data were first log-transformed and then standardized across all trials.

All other data were analyzed using MATLAB R2016b with FieldTrip (Oostenveld et al., 2011) (RRID: SCR_004849), the MEG and EEG Toolbox Hamburg (METH, Guido Nolte; RRID: SCR_016104), or custom made scripts. The physiological continuous recording of each session was first cut into epochs of variable length. Each trial was cut 1,250 ms earlier to stimulus movement onset and 1,250 ms after offset of the reappeared stimulus. Trial length therefore varied between 4,717 and 6,183 ms. To prevent that the timing in a given trial was not exactly as intended, e.g., by short movement interruptions of the stimulus, we removed trials which contained MEG marker timings that differed from the intended timing of the moving stimulus in the trial by at least one frame (17 ms). Thus, we excluded on average 1.2 trials in each participant and each session (range: 0 – 24 trials).

Moreover, trials containing strong muscle artifacts or jumps were detected by semiautomatic procedures implemented in FieldTrip and excluded from analysis. The remaining trials were filtered with a high-pass filter at 0.5 Hz, a low-pass filter at 170 Hz, and three band-stop filters at 49.5–50.5 Hz, 99.5–100.5 Hz and 149.5–150.5 Hz and subsequently down-sampled to 400 Hz.

We performed an independent component analysis (infomax algorithm) to remove components containing eye-movements, muscle, and cardiac artefacts. Components were identified by visual inspection of their time course, variance across samples, power spectrum, and topography. On average, 25.7 ± 8.6 components were rejected in each participant and each session. All trials were again visually inspected and trials containing artefacts that were not detected by the previous steps were removed.

As a final step, using procedures described by Stolk *et al.* (Stolk et al., 2013) and online (http://www.fieldtriptoolbox.org/example/how_to_incorporate_head_movements_in_MEG_analysis/) we identified trials in which the head position of the participant differed by 5 mm from the mean circumcenter of the head position from the whole session (on average: 2.6 trials per participant and session, range: 0 – 86 trials) and excluded them from further analysis. 670.2 ± 26.7 trials of the total of 720 trials remained from pre-processing on average per participant in each session.

### Quantification and statistical analysis

In the current experiment, we introduced a control condition that was physically identical to our temporal prediction tasks (until reappearance in the tactile condition) in order to account for processes that are not directly related temporal predictions. Hence, for most of our statistical analyses, we were interested in comparing the two temporal prediction tasks with the luminance matching control task, respectively, and not in comparing the two temporal prediction tasks with each other. Therefore, instead of computing an analysis of variance across all three conditions, we directly computed two separate *t*-tests for the comparison of the visual or the tactile temporal prediction with the luminance matching task, respectively, and accounted for multiple comparisons by adjusting the alpha level.

#### Psychometric curve

We did not provide participants with feedback about the correctness of their response. Hence, participants responded within their individual framework of a “subjectively correct” reappearance timing or a “subjectively equal” luminance of the stimulus, respectively. To obtain these subjective points of “right-on-time” (ROT) in the temporal prediction tasks or the “points of subjective equality” (PSE) in the luminance matching task, we fitted a psychometric curve to the behavioral data of each participant from all trials in each condition. First, for each timing difference or luminance difference, respectively, we computed the proportion of “too late” or “brighter” answers for each participant. Then, we fitted a binomial logistic regression (psychometric curve) using the glmfit.m and gmlval.m functions provided in MATLAB. The fitted timing or luminance difference value at 50% proportion “too late” or “brighter” answers was determined as ROT or PSE for each participant, respectively. To test for a significant bias towards one of the answers, we tested the ROT or PSE from all participants against zero using one-sample *t*-tests (α = .05 / 3 = .017). The steepness of the psychometric function was computed as the reciprocal of the difference between fitted timing or luminance difference values at 75% and 25% proportion “too late” or “brighter” answers, respectively.

#### Mixed regression model for reaction times

To test whether reaction times were dependent on the timing difference of the reappearing stimulus, we fitted a random intercept and slope mixed model to reaction times from all trials using the categorial variable *condition* (with the luminance matching task as reference level) and *timing difference* as well as their interaction as fixed effects. Since in the temporal prediction conditions we expected reaction times to be slowest for timing differences around zero and faster for high timing differences, we used a second-order polynomial term for *timing differences.* Subject ID was used as grouping variable to model an individual intercept for each participant, and *timing difference* was modeled with random slope. We used R including the *lme4* package for computing the mixed-effect model, and the package *parameters* to compute p-values using the “Kenward” option, which estimates p-values for fixed effects using the Kenward-Roger approach (Kenward and Roger, 1997).

#### Spectral power

We decomposed the MEG recordings into time-frequency representations by convolving the data with complex Morlet’s wavelets (Cohen, 2014). The recording of each trial and channel was convolved with 40 complex wavelets, logarithmically spaced between 0.5 to 100 Hz. With increasing frequency, the number of cycles for each wavelet logarithmically increased from two to ten cycles. For all analyses of the MEG data, we considered subjectively correct trials only, i.e., trials in which participants answered correctly based on their individual ROT. To correct for trial count differences between the tasks, we stratified the number of trials for each participant for the three different conditions by randomly selecting as many trials for each condition as the number available from the condition with lowest trial count.

Since the temporal dependencies between the movement onset, disappearance behind the occluder and reappearance of the stimulus varied strongly between trials, averaging across trials would heavily smear the power estimates of the different stages within each trial. To obtain an estimate of spectral power modulations related to the different events in our experimental paradigm, we cut each trial further into four separate, partly overlapping windows (see Figure 2A): a “Baseline” window from −550 to −50 ms earlier to movement onset; a “Movement” window from −50 to 950 ms relative to the movement onset; a “Disappearance” window from −350 to 950 ms relative to complete disappearance of the stimulus behind the occluder; and a “Reappearance” window from −350 to 450 ms relative to the (first frame) reappearance of the stimulus. Spectral power estimates were then averaged across all trials belonging to the same condition in each window and binned into time windows of 100 ms (centered on each full deci-second). All power estimates were normalized using the pre-stimulus baseline window from −500 to −200 ms earlier to movement onset.

For all statistical analyses on sensor level, we first flipped all sensors of participants, who saw the stimulus moving from right to left, at the sagittal midline, i.e., the anterior-posterior axis. This made sure that lateralized activity due to the lateralized stimulation was comparable across groups. From this on, we considered all participants as if for everyone the stimulus was moving from the left to the right side. Channels that did not have a counterpart on the opposite site were excluded from further analyses. In order to obtain an overview of the spectral power modulations related to the different events within the trials, we then averaged the power estimates across all channels and conditions (grand average) and tested each timefrequency pair of the Movement, Disappearance and Reappearance windows against the prestimulus baseline using paired-sample *t*-tests. We controlled for multiple comparisons by employing cluster-based permutation statistics as implemented in FieldTrip(Maris and Oostenveld, 2007). In this procedure, neighboring time-frequency bins with an uncorrected *p*-value below 0.05 are combined into clusters, for which the sum of t-values is computed. A null-distribution is created through permutations of data across participants (*n* = 1,000 permutations), which defines the maximum cluster-level test statistics and corrected p-values for each cluster. For each window, a separate cluster-permutation test was performed (*α* = .05; liberally chosen to observe all ongoing power modulations; see Results section).

Since we were most interested in differences between the conditions during the disappearance time, we subsequently compared the spectral power estimates averaged within the beta range (13–30 Hz; see Results section) at each time point within the disappearance window and all channels from the visual or tactile temporal prediction task with the luminance matching task. We again employed cluster-permutation statistics, this time by clustering neighboring channels and time points. We used a one-sided *α* = .025 / 2 = .0125, since negative and positive clusters were tested separately, and to adjust for the two separate comparisons between the conditions (used throughout the study unless stated differently).

To estimate spectral power in source space, we computed separate leadfields for each recording session and participant based on each participant’s mean head position in each session and individual magnetic resonance images. We used the single-shell volume conductor model (Nolte, 2003) with a 5,003 voxel grid that was aligned to the MNI152 template brain (Montreal Neurological Institute, MNI; http://www.mni.mcgill.ca) as implemented in the METH toolbox. Cross-spectral density (CSD) matrices were computed from the complex wavelet convolved data in steps of 100 ms in the same time windows as outlined above. To avoid biases in source projection, common adaptive linear spatial filters (DICS beamformer (Gross et al., 2001)) pointing into the direction of maximal variance were computed from CSD matrices averaged across all time bins and conditions for each frequency.

All time-frequency resolved CSD matrices were then multiplied with the spatial filters to estimate spectral power in each of the 5,003 voxels and normalized with the pre-stimulus baseline window. Analogous to sensor space, we first flipped all voxels at the y-axis (anterior-posterior axis) for the half of participants that saw the stimulus moving from right to left earlier to further statistical analysis. We then averaged across all time bins within the disappearance window and utilized cluster-based permutation statistics to identify clusters of voxels that show statistical difference in beta power between each of the temporal prediction tasks and the luminance matching task.

#### Inter-trial phase consistency

We computed ITPC estimates from the complex time-frequency representations obtained from the wavelet convolution as described in the *Spectral power* section above. In each time sample and trial, the phase of the complex data was extracted (using the function angle.m in MATLAB). ITPC was then computed across all subjectively correct and stratified trials within each of the four time windows in all frequencies as

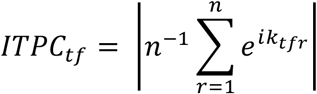

where *n* is the number of trials and *k* the phase angle in trial *r* at time-frequency point *tf* (Cohen, 2014). In other words, ITPC is the length of the mean vector from all phase vectors with length 1 across all trials at a given time-frequency point. Values for ITPC can vary between 0 and 1, where 0 means that at a given time-frequency point there is no phase consistency across trials at all and 1 means all trials show the exact same phase. Similar to spectral power, we averaged ITPC estimates again in bins of 100 ms and plotted all time windows averaged across all channels and conditions to obtain a general overview of ITPC estimates at all events during the trial.

Since we were most interested in ITPC related to stimulus disappearance behind the occluder, we subsequently computed ITPC in a longer time window from −1,900 ms to 1,900 ms centered around time of complete stimulus disappearance behind the occluder. Thus, we took advantage of the fact that the onset of other events within each trial, such as the movement onset and the reappearance of the stimulus, strongly jittered across all trials and strong contributions of these events to ITPC could thereby be reduced (see Fig. S3). For statistical analysis, we first averaged ITPC estimates within a frequency band of 0.5 to 3 Hz (see Results) and then computed cluster-based permutation statistics across all 100 ms time bins within the 3,800 ms long window and all sensors between each of the temporal prediction tasks and the luminance matching task.

ITPC on source level was computed using the same leadfields and common beamformer filters as for spectral power (see above). The complex time-frequency representations obtained from the wavelet convolution within the 3,800 ms long window on sensor level were multiplied with the filters to obtain the time-frequency representations in each of the 5,003 voxels. ITPC was computed for each time sample and frequency and then averaged within the time window showing statistically significant difference between the temporal prediction tasks and the luminance matching task on sensor level and within the pre-defined frequency band of 0.5 to 3 Hz. Cluster-based permutation statistics were employed to identify clusters of voxels showing statistically significant differences in ITPC between the conditions on source level.

Correlations between condition-wise source level ITPC estimates and the steepness of each individual’s psychometric function were computed using Pearson correlations in each of the 5,003 voxels within the grid. For this analysis, we averaged ITPC estimates from time bins of 0 to 1,000 ms with respect to the disappearance of the stimulus within the pre-defined delta band of 0.5 to 3 Hz. Multiple comparisons were accounted for by using cluster-based permutation statistics as implemented in FieldTrip (*α* = .025 / 3 = .008)

#### Delta power control analyses and mixed models

For control analyses of delta power differences between the conditions, we computed delta power using the same wavelet convolution approach as described for ITPC for the enlarged time windows between −1,900 ms to 1,900 locked to stimulus disappearance. To obtain total delta power, we computed power in each single trial first and then averaged power within the delta band (0.5 – 3 Hz) and the respective channels showing the strongest ITPC effect (see Fig. 3D) for each time bin and condition. Induced power was obtained by first averaging all trials in each condition and channel in the time domain, i.e., by computing an ERF in each channel and condition, and then subtracting this average from all single trials in each channel and condition separately. After subtracting the ERF, power was estimated as described for total power above. Delta power of the ERF itself was estimated by applying a wavelet convolution to the ERF, i.e., the average across trials, in each condition and channel and subsequently averaging power estimates within the delta band and the respective channels. All time courses were baseline corrected with a pre-disappearance window of – 1,500 to −500 ms relative to disappearance in each condition.

To further examine the effect of delta power on ITPC, we computed random intercept and random slope mixed-effects models using *condition* and *time* as well as their interaction as fixed effects for predicting ITPC. One model also included delta power as an additional covariate, the other one did not. We first averaged delta ITPC as well as delta power (0.5 – 3 Hz) from each condition and each time bin (−1.900 ms – 1.900 ms) within the channels showing the strongest effect for ITPC (see Fig. 3D). ITPC as well as the baseline-corrected power values were standardized across all data for an easier interpretation of the model estimates. *Subject ID* was used as grouping variable to model an individual intercept for each participant, and *time* was modeled as random slope. To ensure a flexible relationship between time and ITPC, we modeled *time* using natural cubic splines with 10 degrees of freedom. For plotting, we computed ITPC values as predicted by the interaction between *condition* and *time* and back-transformed the values to the original scale for an easier evaluation. As for the reaction time model, we used R including the *lme4* package for computing the mixed-effect model, the package *parameters* to compute p-values using the “Kenward” option, as well as the package *splines* for generating the natural cubic splines.

## Acknowledgements

We thank Florian Göschl, Tessa Rusch, Marina Fiene and Guido Nolte for valuable discussions. This work was funded by grants from the DFG (SFB TRR 169/B1 and SFB 936/A3 to A.K.E.).

## Author contributions

Conceptualization, J.D., A.K.E, P.W., A.M.; Methodology, J.D., A.K.E.; Software, J.D.; Formal Analysis, J.D.; Investigation, J.D.; Writing – Original Draft, J.D.; Writing – Review & Editing, A.K.E., P.W., A.M., D.Z.; Visualization, J.D.; Funding Acquisition, A.K.E.; Supervision, A.K.E.; Project Administration, A.K.E., D.Z.; Resources, A.K.E.

## Data availability

Custom scripts and data will be made available upon full submission of the manuscript.

