## supplemental information for "Delta phase resets mediate non-rhythmic temporal prediction"

### Supplementary Results

| Predictors | Estimates | t | p |
|---|---|---|---|
| (Intercept) | 0.13 | 1.38 | 0.180 |
| Condition: Visual prediction | -0.03 | -1.34 | 0.179 |
| Condition: Tactile prediction | -0.26 | -14.21 | **<0.001** |
| Timing difference | -0.04 | -2.48 | **0.019** |
| Timing difference$^2$ | 0.02 | 2.42 | **0.016** |
| Visual prediction : timing difference | -0.13 | -10.53 | **<0.001** |
| Tactile prediction : timing difference | -0.02 | -1.25 | 0.212 |
| Visual prediction : timing difference$^2$ | -0.11 | -8.18 | **<0.001** |
| Tactile prediction : timing difference$^2$ | -0.10 | -7.52 | **<0.001** |
| **Random effects** | | | |
| $\sigma^2$ | 0.40 | | |
| $\tau_{00\ subj}$ | 0.19 | | |
| $\tau_{11\ subj.timing\ difference}$ | 0.00 | | |
| $\rho_{01\ subj}$ | 0.51 | | |
| ICC | 0.33 | | |
| $N_{subj}$ | 23 | | |
| Observations | 14946 | | |
| Marginal $R^2$ / Conditional $R^2$ | 0.058 / 0.365 | | |

**Table S1. Results from random intercept and random slope mixed-effects model for reaction times**. The luminance matching condition was set as reference level in the categorial variable *condition.* P values were computed using the Kenward-Roger approach (see Methods).

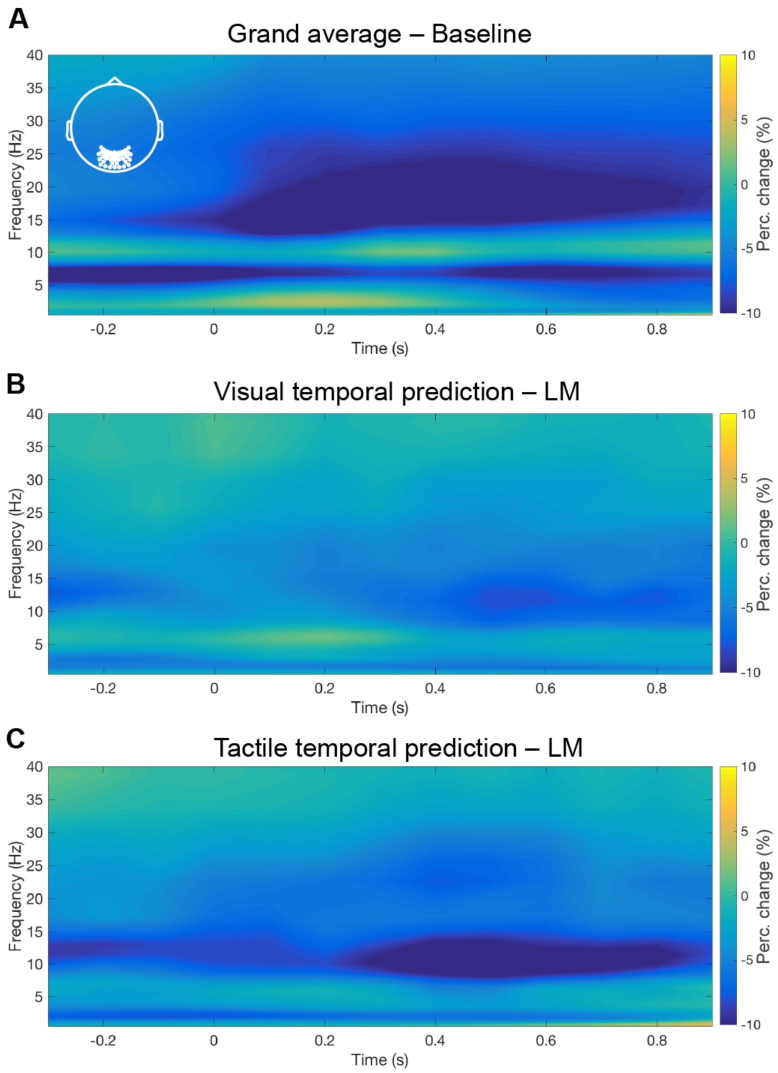


**Figure S1. Low-frequency power modulations averaged across occipital sensors.** (**A**) Power modulations averaged across all three conditions as compared to the pre-stimulus baseline. Cluster-based permutation statistics revealed no significant increase in low-frequency power. Time 0 refers to full disappearance behind the occluder. (**B,C**) Differences in low-frequency power between (**B**) the visual or (**C**) the tactile prediction task and the luminance matching task, respectively, across the same occipital sensors. Especially in these difference plots, it becomes clear that delta power was not stronger during temporal predictions as compared to the luminance matching task. Even when averaging within the 0.5 to 3 Hz delta band, where ITPC differences between the conditions were found, no clusters of significant sensors were found for delta power differences between the conditions.

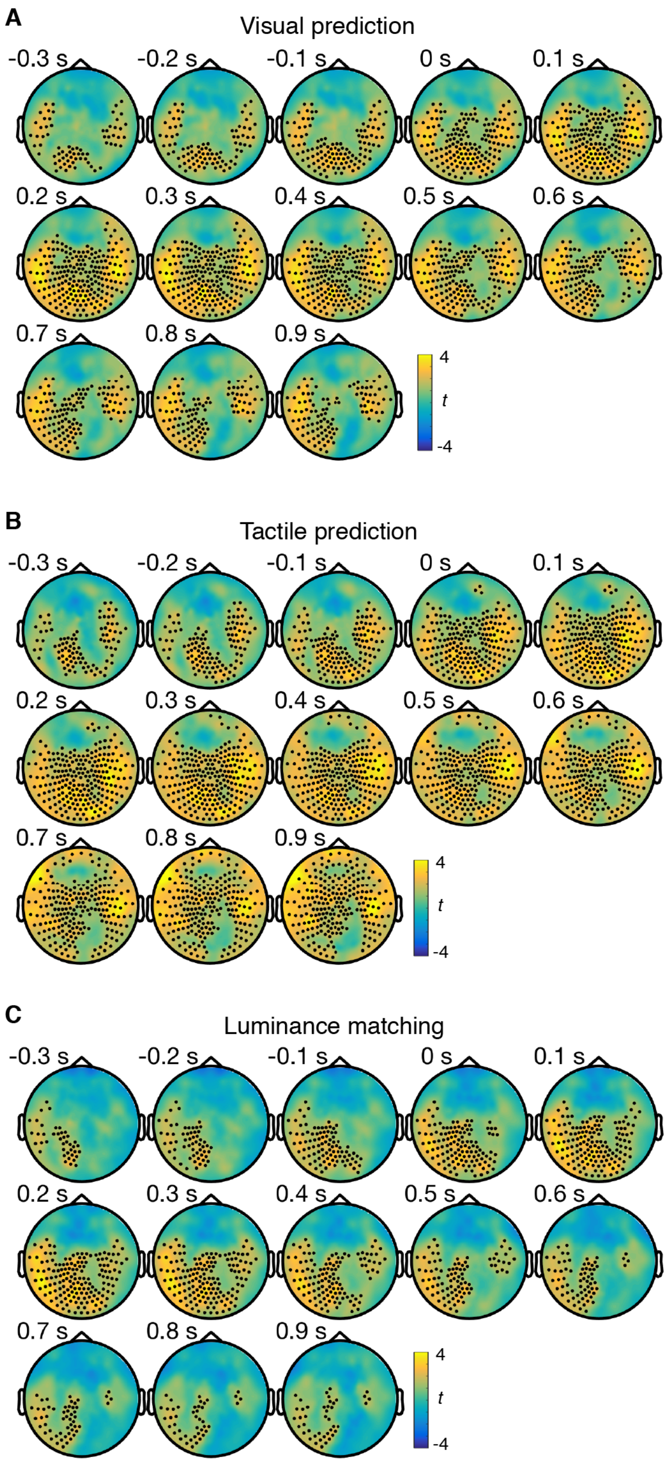


**Figure S2. Condition-specific ITPC differences in the delta band (0.5 – 3 Hz) as compared to pre-stimulus baseline.** Time 0 refers to complete disappearance behind the occluder. In all three conditions, i.e. also the luminance matching condition, ITPC estimates were increased also in posterior sensors (all cluster-$p < .001$).

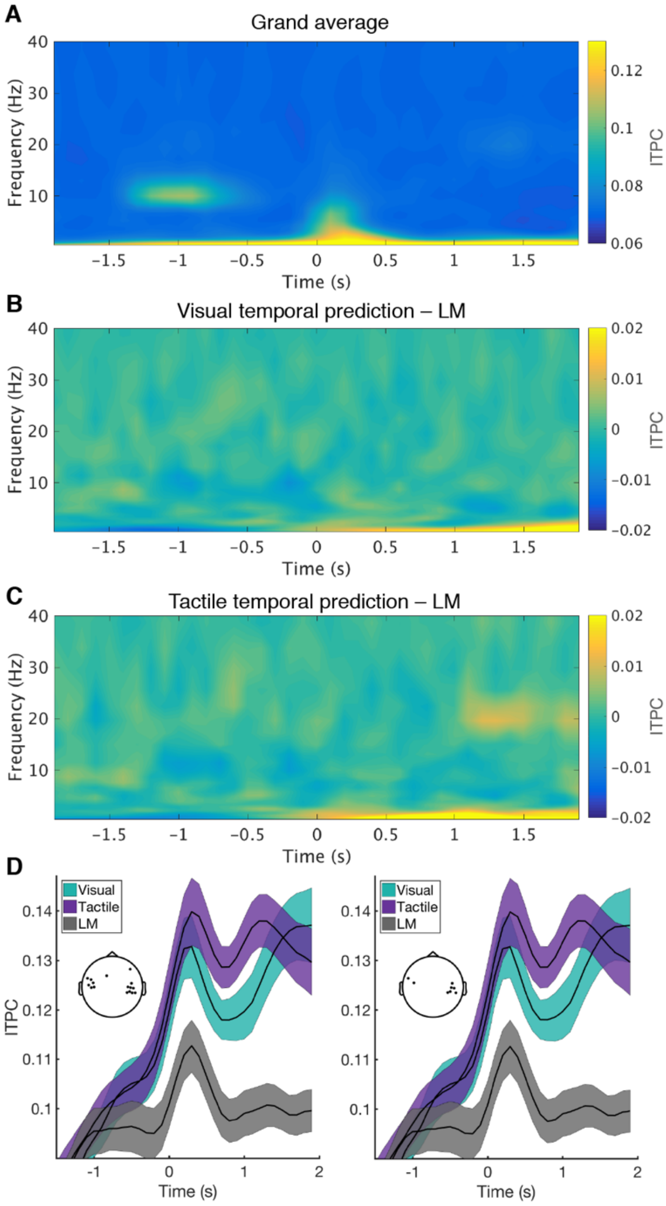


**Figure S3. ITPC estimates in the enlarged analysis window of -1,900 to 1,900 ms.** (**A**) ITPC estimates as an average across all three conditions and all sensors. Note that when centered on stimulus disappearance, the stimulus onset events during *Movement* (between -1,500 and -1,000 ms) and *Reappearance* (roughly between 1,000 and 1,900 ms) strongly jittered across trials and therefore did not affect ITPC estimates in this analysis. At around disappearance, ITPC estimates increased at low frequencies. As compared to the luminance matching task, ITPC was stronger during (**B**) visual predictions as well as (**C**) tactile predictions after disappearance of the stimulus behind the occluder in the delta band. (**D**) Time course of absolute ITPC estimates in each condition averaged across channels showing the top 10% (left panel) or top 5% (right panel) of t values for the comparison of visual temporal prediction with luminance matching (see Figure 3B).

| | *Without delta power* | | | *Adjusted for delta power* | | |
|---|---|---|---|---|---|---|
| **Predictors** | **Estimates** | **t** | **p** | **Estimates** | **t** | **p** |
| (Intercept) | -1.08 | -6.65 | **<0.001** | 0.09 | 0.47 | 0.637 |
| Condition: Visual prediction | -0.10 | -0.68 | 0.497 | -0.10 | -0.72 | 0.472 |
| Condition: Tactile prediction | -0.09 | -0.64 | 0.524 | -0.09 | -0.63 | 0.530 |
| Delta power (z) | | | | 0.38 | 13.41 | **<0.001** |
| time_1 | 0.61 | 3.92 | **<0.001** | -0.59 | -3.35 | **0.001** |
| time_2 | 0.86 | 4.57 | **<0.001** | -0.56 | -2.68 | **0.007** |
| time_3 | 0.57 | 3.11 | **0.002** | -0.64 | -3.25 | **0.001** |
| time_4 | 0.89 | 4.45 | **<0.001** | -0.33 | -1.56 | 0.119 |
| time_5 | 1.54 | 7.31 | **<0.001** | 0.35 | 1.60 | 0.111 |
| time_6 | 0.35 | 1.56 | 0.123 | -0.87 | -3.72 | **<0.001** |
| time_7 | 1.08 | 4.40 | **<0.001** | -0.11 | -0.43 | 0.666 |
| time_8 | 0.77 | 3.21 | **0.002** | -0.43 | -1.80 | 0.073 |
| time_9 | 0.97 | 2.83 | **0.006** | -0.62 | -1.81 | 0.071 |
| time_10 | 0.66 | 2.80 | **0.008** | 1.10 | 4.93 | **<0.001** |
| Visual pred. : time_1 | -0.19 | -0.88 | 0.379 | -0.22 | -1.08 | 0.282 |
| Visual pred. : time_2 | 0.14 | 0.57 | 0.569 | 0.18 | 0.73 | 0.464 |
| Visual pred. : time_3 | 0.34 | 1.50 | 0.134 | 0.37 | 1.65 | 0.099 |
| Visual pred. : time_4 | 0.72 | 2.99 | **0.003** | 0.73 | 3.13 | **0.002** |
| Visual pred. : time_5 | 0.84 | 3.59 | **<0.001** | 0.80 | 3.51 | **<0.001** |
| Visual pred. : time_6 | 0.64 | 2.70 | **0.007** | 0.62 | 2.67 | **0.008** |
| Visual pred. : time_7 | 0.94 | 3.90 | **<0.001** | 0.88 | 3.78 | **<0.001** |
| Visual pred. : time_8 | 1.38 | 6.72 | **<0.001** | 1.26 | 6.33 | **<0.001** |
| Visual pred. : time_9 | 1.39 | 3.81 | **<0.001** | 1.35 | 3.79 | **<0.001** |
| Visual pred. : time_10 | 1.41 | 8.22 | **<0.001** | 1.42 | 8.52 | **<0.001** |
| Tactile pred. : time_1 | -0.06 | -0.26 | 0.791 | -0.11 | -0.54 | 0.589 |
| Tactile pred. : time_2 | 0.10 | 0.38 | 0.702 | 0.12 | 0.48 | 0.629 |
| Tactile pred. : time_3 | 0.25 | 1.09 | 0.276 | 0.29 | 1.32 | 0.186 |
| Tactile pred. : time_4 | 0.90 | 3.76 | **<0.001** | 0.91 | 3.91 | **<0.001** |
| Tactile pred. : time_5 | 1.04 | 4.42 | **<0.001** | 0.99 | 4.38 | **<0.001** |
| Tactile pred. : time_6 | 1.17 | 4.90 | **<0.001** | 1.09 | 4.69 | **<0.001** |
| Tactile pred. : time_7 | 1.50 | 6.21 | **<0.001** | 0.96 | 4.03 | **<0.001** |
| Tactile pred. : time_8 | 1.11 | 5.42 | **<0.001** | 0.41 | 1.99 | **0.047** |
| Tactile pred. : time_9 | 1.27 | 3.46 | **0.001** | 0.82 | 2.31 | **0.021** |
| Tactile pred. : time_10 | 1.18 | 6.87 | **<0.001** | 0.86 | 5.10 | **<0.001** |
| **Random effects** | | | | | | |
| $\sigma^2$ | 0.33 | | | 0.31 | | |
| $\tau_{00}$ | 0.15 $_{subj}$ | | | 0.13 $_{subj}$ | | |
| $\tau_{11}$ | 0.07 $_{subj.time}$ | | | 0.06 $_{subj.time}$ | | |
| $\rho_{01}$ | 0.11 $_{subj}$ | | | -0.06 $_{subj}$ | | |
| ICC | 0.31 | | | 0.30 | | |
| N | 23 $_{subj}$ | | | 23 $_{subj}$ | | |
| Observations | 2691 | | | 2691 | | |
| Marginal $R^2$ / Conditional $R^2$ | 0.485 / 0.644 | | | 0.519 / 0.666 | | |

**Table S2. Results from the random intercept and random slope mixed-effects models for ITPC**. The luminance matching condition was set as reference level in the categorial variable *condition.* P-values were computed using the Kenward-Roger approach. ITPC and baseline-corrected power values were standardized for an easier interpretation of the estimates.
